# A reference panel for linkage disequilibrium and genotype imputation using whole-genome sequencing data from 2,680 participants across India

**DOI:** 10.1101/2025.06.30.662450

**Authors:** Zheng Li, Wei Zhao, Xiang Zhou, Yuk Yee Leung, Gerard D. Schellenberg, Li-San Wang, Sharmistha Dey, Jinkook Lee, Jennifer A. Smith, Aparajit B. Dey, Sharon L.R. Kardia

**Author notes:** Correspondence to Wei Zhao and Jennifer A. Smith. These authors contribute equally to this manuscript (Zheng Li, Wei Zhao). Senior author (Jennifer A. Smith, Aparajit B. Dey, Sharon L.R. Kardia).

## Abstract

India is the most populous country globally, yet genetic studies involving Indian individuals remain limited. The Indian population is composed of many founder groups and has a mixed genetic ancestry, including an ancestral component not observed anywhere outside of India. This presents a unique opportunity to uncover novel disease variants and develop more tailored medical interventions. To facilitate genetic research in India, a crucial first step is to create a foundational resource that serves as a benchmark for future population studies and methods development. To this end, we have constructed the largest and most nationally representative linkage disequilibrium and genotype imputation reference panels in India to date, using high-coverage whole-genome sequencing data of 2,680 Indian participants from the Longitudinal Aging Study in India-Harmonized Diagnostic Assessment of Dementia (LASI-DAD). As an LD reference panel, LASI-DAD includes 69.5 million variants, representing 170% and 213% increases relative to the 1000 Genomes Project (1000G) and TOP-LD panels, respectively. Besides serving as an LD lookup panel, LASI-DAD facilitates various statistical analyses that rely on precise LD estimates. In a polygenic risk score (PRS) analysis, LASI-DAD improved the predictive performance of PRS by 2.1% to 35.1% across traits and studies. As an imputation reference panel, LASI-DAD improved the imputation accuracy by 3% to 101% (mean = 38%) compared to the TOPMed panel (Version R3) and by 3% to 73% (mean = 27%) compared to the Genome Asia Pilot (GAsP) panel across different minor allele frequencies. The LASI-DAD reference panel is publicly available to benefit future studies.

## Introduction

India, with a population of over 1.4 billion, is home to more than 4,500 anthropologically defined groups that are characterized by a rich diversity in languages, religions, and cultures.^1^ The Indian population largely descends from a mixture of two genetically divergent populations: Ancestral North Indians (ANI), who are related to Central Asians, Middle Easterners, Caucasians, and Europeans; and Ancestral South Indians (ASI), who are not related to any groups outside the subcontinent and are only distantly related to the indigenous Andamanese Islanders.^2, 3^ Due to this unique ASI component, the Indian population has a distinctive genetic structure and harbors many unique variants not present elsewhere. Thus, genetic variants identified from populations in other areas of the world may not have the same effects in Indians, and many variants specific to the Indian population could play significant roles in predisposing this population to certain diseases. Furthermore, due to consanguineous and endogamous marriages, the ancestral Indian population experienced founder events that are more extreme than Ashkenazi Jews and Finnish populations, which led to many genetically diverse subgroups.^4^ As a result, the frequency of deleterious or protective variants may increase substantially, leading to increased power for gene mapping. All of these highlight the critical need and potential for new discovery of variants (otherwise rare outside India) from genetic studies in this population. Despite the need and opportunity, India has been largely underrepresented in genetic studies, making up less than 2% of human study participants in the catalog of genome-wide association studies (GWAS catalog).^5, 6^ This lack of representation hinders the discovery of novel disease variants and the development of more tailored medical interventions or treatments towards the Indian population.

To facilitate genetic research in India, a crucial first step is to create an Indian-representative reference panel that can serve as a resource for future studies in India as well as methods development. This resource may serve as a reference panel for linkage disequilibrium (LD) and genotype imputation for the Indian population. LD refers to the non-random co-occurrence of alleles at different genomic loci.^7^ A robust and representative estimate of genome-wide LD statistics for a given population is critical for a wide range of genetic analyses,^8–12^ such as statistical fine-mapping and polygenic risk score (PRS) construction. Genotype imputation, on the other hand, estimates unobserved genotypes in samples with sparse microarray data and has been widely used in GWAS to boost power, fine-map candidate causal variants, and facilitate meta-analysis across studies.^13^ Extensive efforts have been made by several initiatives such as the 1000 Genomes Project (1000G), Trans-Omics for Precision Medicine (TOPMed), and Genome Asia Pilot (GAsP) project to establish reference panels.^14–17^ However, these reference panels have included a limited number of Indian samples and/or primarily focused on expatriate communities outside of India. For example, 1000G includes 489 unrelated individuals from South Asia, which includes only 103 Gujarati Indians in Houston, Texas and 102 Indian Telugu individuals in the United Kingdom. Likewise, the only South Asian representation in TOPMed comes from the Pakistani Risk of Myocardial Infarction Study (PROMIS) with a sample size of 644. While GAsP consists of a broad range of populations across Asia with a total sample size of 1,739, only 598 individuals are from India. This limited sample size and representation has restricted the utility of these resources to serve as reference panels for the Indian population.

In the present study, we have constructed the largest and most nationally representative LD and genotype imputation reference panel in India to date, using high-coverage (30x) whole genome sequencing (WGS) data from 2,680 Indian participants from the Longitudinal Aging Study in India-Harmonized Diagnostic Assessment of Dementia (LASI-DAD). For the LD reference panel, we characterized LD patterns in LASI-DAD by identifying LD blocks and evaluating regional differences in LD between LASI-DAD and four super-populations from 1000G. Given the genetic heterogeneity in this population, we further characterized and compared LD patterns by sub-populations within LASI-DAD and evaluated how such differences manifest in PRS performance across sub-populations. We further assessed the utility of this LD reference panel in facilitating PRS construction in South Asians. The LASI-DAD LD reference panel covers a substantially higher number of genetic variants in India than previous panels and facilitates various statistical analysis that rely on precise LD estimates. To evaluate LASI-DAD as an imputation reference panel, we performed imputation using phased LASI-DAD WGS data or other widely-used reference panels, including TOPMed and GAsP, and compared the imputation accuracy across the spectrum of minor allele frequencies. As an imputation reference panel, LASI-DAD enables the imputation of many Indian-specific variants that are not present in the other panels with improved imputation accuracy.

## Material and methods

### The Longitudinal Aging Study in India - Harmonized Diagnostic Assessment of Dementia (LASI-DAD)

LASI is a nationally representative survey of more than 70,000 adults aged 45 years and older in India that aims to understand the health, economic, and social well-being of the older adults in India. LASI-DAD is an ancillary study of LASI that subsampled adults who were at least 60 years of age from 18 states and union territories across India to specifically focus on late-life cognition and dementia. Participants were selected based on a two-stage stratified sampling approach that oversampled individuals with a high risk of cognitive impairment.^18^ Briefly, participants in the larger LASI cohort were first classified into those at high risk or low risk of cognitive impairment based on cognitive testing or on the proxy report for those who did not complete the cognitive tests. Finally, an approximately equal number of participants were randomly sampled from the high risk and low risk strata with a target sample size for each state/territory proportional to the population size.

A total of 2,762 LASI-DAD participants who consented to blood sample collection were carried forward for whole-genome sequencing (WGS) at MedGenome, Inc. (Bangalore, India) at an average read depth of 30x. Genotype calling and quality control (QC) of the raw WGS data were conducted at the Genome Center for Alzheimer’s Disease (GCAD) at the University of Pennsylvania. Specifically, samples were excluded due to low sequencing coverage, sample contamination, sex mismatch, discordance with previous genotyping array data,^19^ unexpected duplications, or serving as technical controls. A total of 2,680 samples were retained in the analysis after sample-level QC. The number of samples by state/territory is reported in Table S1, with states/territories grouped by geographic regions. For genotype-level QC, each genotype was examined and set to missing if its read depth (DP) was less than 10 or genotype quality (GQ) score was less than 20. For variant-level QC, a variant was excluded if it was monomorphic, fell in the 99.8% Variant Quality Score Recalibration (VQSR) Tranche or higher, had a call rate less than or equal to 80%, or had an average depth greater than 500 reads. We further excluded variants in low-complexity regions as identified by mdust.^20^ After variant-level QC, a total of 71,109,961 biallelic variants that included 66,204,161 autosomal single nucleotide polymorphisms (SNPs) and 4,905,800 indels were retained in the analysis. We phased the genotype data with Eagle2.4^21, 22^ without a reference panel following the user manual of Eagle2.4 (https://alkesgroup.broadinstitute.org/Eagle/#x1-130004). Finally, we estimated genetic principal components (PCs) using PC-AiR in both LASI-DAD and the combined samples of LASI-DAD and 1000G as described previously.^23, 24^

### Characterizing and comparing linkage disequilibrium patterns between LASI-DAD and 1000 Genomes Project super-populations

We characterized LD patterns by identifying LD blocks and evaluating the variation in LD (varLD) scores. For both LD block identification and varLD score calculation, we focused on a total of 2,607 unrelated samples based on a kinship coefficient threshold of 2^-3.5^ (or 0.088) obtained from PC-Relate.^25^ LD blocks represent segments of the genome where genetic variants are often inherited together. The identification of LD blocks can be useful in many ways for genetic studies. For example, some polygenic risk score methods estimate SNP effects one block at a time to achieve a good balance between modeling accuracy and computational efficiency.^26–28^ As another example, local genetic correlation analysis quantifies the genetic similarity of complex traits within individual LD blocks.^29–31^

We applied LDetect^32^ to identify approximately independent LD blocks in LASI-DAD, and compared with those identified in four super-populations from 1000G: African (AFR), East Asian (EAS), European (EUR), and South Asian (SAS). LDetect first partitions each chromosome into smaller chunks to facilitate parallel processing. For each chunk, it computes a covariance matrix between all pairs of SNPs using the shrinkage estimator from Wen and Stephens.^33^ Notably, both steps require the input of a recombination map to refine the boundary of chunks and to adjust the level of shrinkage. Following MacDonald et al.,^34^ we used the recombination map inferred for Gujarati Indians in Houston (n = 113) in the 1000G phase 3 data.^35^ Recombination rates were linearly interpolated to all SNPs in the LASI-DAD data. Afterward, LDetect converts the covariance matrix into a vector with each element representing the sum of elements along a corresponding antidiagonal of the matrix. Boundaries of LD blocks appear as “dips” in the vector and can be identified by first smoothing the vector and then searching for local minima. In the analysis, we focused on SNPs with MAF ≥ 1% and set the average block size to 7,000 SNPs following MacDonald et al.^34^ To compare LASI-DAD and the 1000G super-populations, we directly obtained LD blocks from MacDonald et al.^34^ Notably, the American (AMR) super-population was not included in MacDonald et al.,^34^ likely due to its extensive genetic admixture.

While LDetect is effective in capturing large LD blocks, it is not well-suited for identifying finer LD structures within each block, which can provide additional insights into local LD variation. To further identify LD blocks at a finer scale, we applied Big-LD.^36^ Big-LD proceeds by first identifying LD clusters through a modified clique-based clustering (CLQ) algorithm. Each cluster consists of SNPs that are in strong LD with each other yet are not necessarily physically consecutive. Big-LD then expands each LD cluster to form a consecutive genomic segment that includes all SNPs inside the region. Finally, by merging mutually overlapping segments based on interval graph modelling, Big-LD effectively partitions the genome into a set of finer LD blocks. Big-LD was implemented in the gpart R package (version 1.17.0). We set the |r| threshold to be its default value of 0.5 in the CLQ algorithm, where |r| represents the absolute correlation coefficient between the dosage values of a SNP pair. For the comparison of LD blocks at the fine scale, we applied Big-LD to the same four super-populations in 1000G. Specifically, we obtained the genotype data for 3,202 samples from the 1000G 30x dataset on GRCh38 (https://www.internationalgenome.org/data-portal/data-collection/30x-grch38) and restricted the samples to 2,504 unrelated individuals as reported by 1000G for our analysis.^15^ Following MacDonald et al.,^34^ we excluded the African Caribbean in Barbados (ACB; n = 96) and African Ancestry in Southwest US (ASW; n = 61) from the AFR population due to genetic admixture, as well as the Finnish in Finland (FIN; n = 99) from the EUR population due to genetic isolation. The total number of samples included were 504 for AFR, 504 for EAS, 404 for EUR, and 489 for SAS.

In addition to LD block identification, we compared LD patterns by evaluating the varLD score between LASI-DAD and the four 1000G super-populations.^37^ varLD examines regional patterns of LD between two populations by comparing the signed r^2^ values across all pairs of SNPs in a genomic region with a MAF ≥ 1% that are not strand-ambiguous in both populations, where r^2^ represents the squared correlation coefficient between the dosage values of a SNP pair. Specifically, for a particular genomic region, we first calculated the LD matrix for the two populations, defined as a SNP-by-SNP correlation matrix with each matrix element representing the signed r^2^ value between a SNP pair. We then obtained the eigenvalues of the LD matrix from each population and calculated the varLD score as the sum of the absolute differences between the corresponding eigenvalues derived from the two populations. We employed a sliding-window approach and calculated varLD scores within consecutive windows of 50 SNPs. Each consecutive window was obtained by shifting the current window by one SNP in the direction of the forward strand.^25^

### Characterizing LD patterns by LASI-DAD sub-population

In addition to comparisons with 1000G super-populations, we evaluated LD patterns across three LASI-DAD sub-populations, defined according to the population structure in India. The population structure in India is correlated with geography, with majority of the individuals lying along a North/South cline with varying proportions of Ancestral North Indian (ANI) and Ancestral South Indian (ASI) ancestries. ANI is genetically related to West Eurasians while ASI is related (distantly) to indigenous Andaman Islanders.^2^ In contrast, individuals in the east of India are considered out-of-cline due to admixture with additional ancestral populations such as East Asians. Following Kerdoncuff et al.,^38^ we clustered individuals into those that lie along the North/South cline and those that are out of the cline. For individuals that lie along the North/South cline, we estimated the proportion of ANI ancestry (%ANI) using the f_4_-statistic implemented in the ADMIXTOOLS R package (version 1.3.0).^39^ The f_4_-statistic measures the excess of West Eurasian ancestry in a test individual relative to the Onge population based on an Indian population model described by Moorjani et al.^3^ Finally, we grouped individuals from LASI-DAD into three sub-populations based on their respective cline and %ANI values to facilitate Indian sub-population analysis. The three sub-populations included 945 individuals in the high %ANI group (0.540 ≤ %ANI < 0.824), 957 in the low %ANI group (%ANI ≤ 0.540), and 778 that were out of the North/South cline. We conducted this analysis on the set of unrelated samples, including 935, 941, and 731 individuals in the respective groups. For simplicity, we focused on the evaluation of varLD scores in the sub-population analysis.

### Transferability of polygenic risk scores

A polygenic risk score (PRS) captures an individual’s genetic predisposition to a complex trait or disease and is often calculated as the sum of SNP genotypes weighted by their effect sizes estimated from GWAS. However, most GWASs have been conducted in individuals of European ancestry^40^ and PRS derived from these GWASs may be substantially less predictive in other genetic ancestries.^11^ We evaluated the transferability of the PRS derived from GWASs of primarily European ancestry to LASI-DAD individuals. Specifically, we obtained PRS weights for height and body mass index (BMI) from the Genetic Investigation of Anthropometric Traits (GIANT) consortium. For height, we downloaded PRS weights that were generated by applying SBayesC to a cross-ancestry GWAS meta-analysis^41^ (https://portals.broadinstitute.org/collaboration/giant/index.php/GIANT_consortium_data_files). The meta-analysis included 5,380,080 individuals across 281 studies from the GIANT consortium and 23andMe, with 75.8% of the individuals being primarily of European ancestry and 1.4% being primarily of South Asian ancestry. For BMI, we estimated PRS weights by applying PRS-CS^26^ to a GWAS meta-analysis of GIANT consortium studies and UK Biobank.^42^ The meta-analysis included ∼700,000 individuals of primarily European ancestry. After obtaining the PRS weights, we evaluated the performance of PRS in the three LASI-DAD sub-populations (high %ANI, low %ANI, and out-of-cline) using incremental R^2^. Following Yengo et al.,^41^ we calculated the incremental R^2^ for height by first centering and standardizing height to have a mean of zero and a variance of one within each sex. The incremental R^2^ was then calculated as the difference in the coefficient of determination between two linear regression models: one that includes age and the top 20 genotype PCs as covariates, and the other that additionally includes the height PRS. For BMI, we followed a similar procedure to calculate the incremental R^2^.

### Providing LASI-DAD as a reference panel for linkage disequilibrium

A robust and representative estimate of genome-wide LD statistics for a given population plays an essential role for a wide range of genetic analyses.^8–12^ As the largest and most nationally representative WGS study of the Indian population, LASI-DAD has the potential to provide more accurate estimates of LD with higher coverage of genetic variants, serving as a valuable resource for LD in this population. We evaluated the utility of LASI-DAD WGS data as an LD reference panel to facilitate both LD lookup and various statistical analyses that rely on precise LD estimates. We constructed an LD reference panel using 2,607 unrelated individuals from the LASI-DAD WGS data and provided it as a public resource to facilitate future research in the community. Specifically, for all pairs of genetic variants within 1 million base pairs (Mb) of each other, we computed the squared Pearson correlation coefficient (r^2^) between the phased haplotypes, D-prime (D’), and the direction of the Pearson correlation coefficient. We reported LD statistics for pairs of variants that have an r^2^ statistic ≥ 0.2. Of note, we did not use a minimum minor allele count to exclude any variants. We compared these LD statistics with those obtained from the South Asian populations in 1000G and TOP-LD.^16^ TOP-LD estimated LD statistics for 239 selected South Asian individuals from TOPMed WGS data.^14^ We obtained the LD statistics for TOP-LD directly from its website (http://topld.genetics.unc.edu/about.php). For 1000G, we computed the LD statistics following procedures similar to those used for LASI-DAD and TOP-LD. Of note, an LD panel for the Genome Asia Pilot (GAsP) is not publically available, thus was not included in this comparison.

In addition to serving as an LD lookup panel, LASI-DAD also facilitates various statistical analyses that rely on precise LD estimates. To simplify data sharing while ensuring privacy protection, many statistical methods can directly make use of GWAS summary statistics as input.^43–46^ These summary statistics typically include marginal z-scores for each SNP and a SNP-SNP correlation matrix. The SNP-SNP correlation matrix is commonly referred to as the LD matrix and can be estimated from a reference panel consisting of individuals with the same genetic ancestry as the GWAS. In principle, the closer the LD estimates from the reference panel match those in the GWAS population, the better the results are likely to be in the statistical analysis. This often requires the reference panel to be sufficiently large and representative of the GWAS population. To demonstrate the utility of LASI-DAD as an LD reference panel, we conducted a PRS analysis where we compared the predictive performance of PRS constructed with either the LASI-DAD or 1000G SAS LD reference panel. We estimated PRS weights using PRS-CS, which takes GWAS summary statistics and an external LD reference panel as inputs. We obtained summary statistics for height and BMI from four studies of South Asian ancestry, including a GWAS of height^41^ and an exome-wide association study (ExWAS) of BMI^47^ from the GIANT consortium, and GWASs of both height and BMI from the Genes & Health.^48^ The GIANT data included 77,890 individuals of primarily South Asian ancestry for height and 29,398 for BMI. The Genes & Health data included 36,317 British Pakistani and British Bangladeshi individuals for height and 34,408 for BMI. The summary statistics included marginal association results for 1,339,059 and 246,158 SNPs for height and BMI from the GIANT consortium, and 24,241,756 and 23,922,099 biallelic variants for height and BMI from Genes & Health. We constructed the LASI-DAD LD reference panel following a procedure similar to that of Ge et al.^26^ Specifically, we focused on the 2,607 unrelated samples and filtered out SNPs that have MAF < 1% or are strand-ambiguous. We restricted the SNPs to those in the HapMap 3 panel to achieve a good balance between computational efficiency and prediction accuracy. For each LD block as identified by LDetect, we computed a SNP-SNP correlation matrix, with each element in the matrix representing the Pearson correlation coefficient between the dosage values of a SNP pair. For each study and LD reference panel, we applied PRS-CS to estimate PRS weights. We evaluated the performance of PRS in the full 2,680 LASI-DAD samples using incremental R^2^, calculated following the same procedure as described above.

### Providing LASI-DAD as a reference panel for genotype imputation

Given the unique population structure of India, LASI-DAD has the potential to serve as a valuable resource for genotype imputation to enhance the analysis of array-genotyped samples, especially those collected from the Indian populations. We evaluated the utility of LASI-DAD WGS data as an imputation reference panel and compared it with the GAsP and TOPMed imputation panels. Specifically, we randomly selected 180 samples, 10 from each Indian state or union territory, from the LASI-DAD dataset and selected 530,845 genetic markers that were present on the Illumina Infinium Global Screening Array-24 v2.0 BeadChip. These 180 samples served as a test dataset to evaluate the imputation accuracy. Following Yu et al.,^49^ we evaluated the imputation accuracy by calculating the aggregated squared correlation coefficient (r^2^), computed using aggRSquare (https://github.com/yukt/aggRSquare) which groups variants within a specified MAF range, stacks their imputed dosages and genotype calls from the sequencing data into two separate vectors, and then calculates the aggregated r^2^ as the squared correlation coefficient between the two vectors. We imputed unobserved genotypes with Minimac4 (v1.0.2) using either LASI-DAD (n = 2,500 after excluding the 180 samples from the test dataset), GAsP (n = 1,654), or TOPMed (n = 97,256) as the reference panel. For GAsP and TOPMed, we carried out the imputation directly using the Michigan Imputation Server and the TOPMed Imputation Sever,^50^ respectively. For LASI-DAD, we followed the same analysis pipeline described on the TOPMed Imputation Server for data quality control and imputation. Briefly, we split each chromosome into chunks of 10 Mb, filtered out chunks with fewer than 3 variants, and excluded monomorphic variants. We then conducted genotype imputation using Minimac4 with the same parameter settings as those used by the imputation sever. Besides the imputation with a single reference panel, we also performed meta-imputation by combining imputation results from LASI-DAD with those from GAsP, TOPMed, or both using MetaMinimac2.^49^ We evaluated the imputation accuracy across different ranges of MAF, where the MAFs were calculated among all 2,680 LASI-DAD samples. To facilitate future research in the community, the LASI-DAD reference panel is now available on the Michigan Imputation sever.

## Results

### Characterizing LD patterns in the Indian population from LASI-DAD

We characterized LD patterns in LASI-DAD by identifying LD blocks and evaluating the variation in LD (varLD) scores.^37^ In total, we identified 1,262 blocks in LASI-DAD, which is comparable to the 1,291 blocks identified in the SAS population from 1000G (Figure 1A). For the other 1000G populations, the number of identified blocks was 1,605, 1,360, and 1,143 for AFR, EUR, and EAS, respectively (Figure 1A). As expected, we observed the largest number of LD blocks in the AFR population as Africans are the oldest population ancestrally and there has been more time for recombination to break down the LD structure.^51^ On average, the length of LD blocks was 2.2 million bases (Mb) in LASI-DAD and 1.7 Mb, 2.1 Mb, 2.2 Mb, and 2.5 Mb in the AFR, EUR, SAS, and EAS populations from 1000G, respectively (Figure 1B).

**Figure 1.**
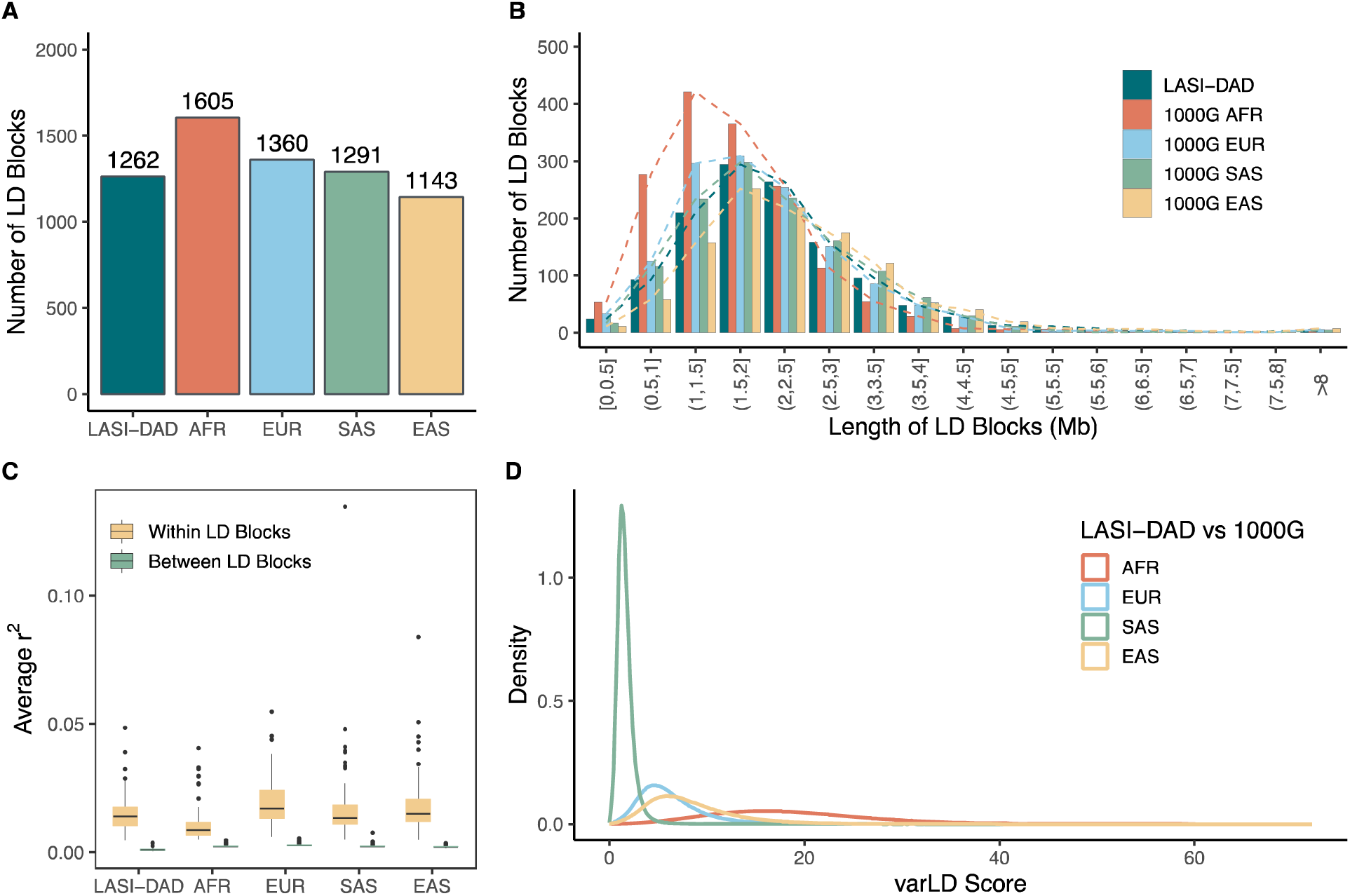
Linkage disequilibrium (LD) patterns in LASI-DAD compared to four super-populations from the 1000 Genomes Project (1000G). The four super-populations include African (AFR), European (EUR), South Asian (SAS), and East Asian (EAS). (**A**) Number of LD blocks identified by LDetect. (**B**) Distribution of LD block lengths in millions of base pairs (Mb), with LD blocks grouped into different length ranges for clear visualization. (**C**) Boxplots showing the distribution of average LD on chromosome 1, calculated either between all pairs of SNPs within each LD block or between SNPs in each LD block and its adjacent two blocks. LD is evaluated as the squared correlation coefficient (r^2^) of genotypes between a SNP pair. (**D**) Distribution of variation in LD (varLD) scores evaluated between LASI-DAD and each of the four super-populations from 1000G.

While the LD blocks identified by LDetect have been widely used by various statistical methods, it is important to note that these blocks represent LD structures at a broad scale, with local LD structures still existing at a finer scale. Indeed, in an evaluation where we compared the average squared Pearson correlation (r^2^) between all pairs of variants within an LD block to the average r^2^ between all pairs of variants in an LD block and its adjacent two blocks, we found that although the average r^2^ was substantially higher within each LD block than with its neighboring two blocks, it remained relatively low, with an average value of 0.015 versus 0.0010 in LASI-DAD (Figure 1C). To better understand the LD structure at a finer scale, we further applied Big-LD.^36^ As expected, the number of LD blocks identified by Big-LD was much larger and ranged from 96,415 to 143,383 across the five populations (Table S2). For these LD blocks, we observed a considerably higher average r^2^ both within and between blocks, with average values of 0.32 and 0.098, respectively, in LASI-DAD (Figure S1). Finally, we visualized an example LD block from LDetect (Figure S2). While this region corresponds to a single LD block at a broad scale, we observed extensive variation both in the local LD structure in this region (Figure S2A) and across different populations (Figure S2B).

In addition to LD block identification, we evaluated regional differences in LD between pairs of populations with varLD scores. As expected, we observed the smallest difference in LD between LASI-DAD and the SAS population from 1000G, with an average varLD score of 1.67 (Figure 1D). This was followed by the EUR (6.21), EAS (8.34), and AFR (18.5) populations.

### Indian population structure and transferability of PRS

In addition to comparisons with 1000G super-populations, we evaluated LD patterns across the three LASI-DAD sub-populations (Figures 2A-B and Table S1). We found that the overall differences in LD across LASI-DAD sub-populations were modest compared to those observed between LASI-DAD and the 1000G EUR, EAS, and AFR populations, but comparable to those between LASI-DAD and the 1000G SAS population (Figure 2C versus Figure 1D). The average varLD scores were 2.10, 1.81, and 1.69 between the out-of-cline and high %ANI groups, high %ANI and low %ANI groups, and the out-of-cline and low %ANI groups, respectively (Figure 2C). This level of difference in LD is consistent with the relative genetic distances in the principal component space, where larger genetic distance corresponds to greater difference in LD (Figures 2B-C).

**Figure 2.**
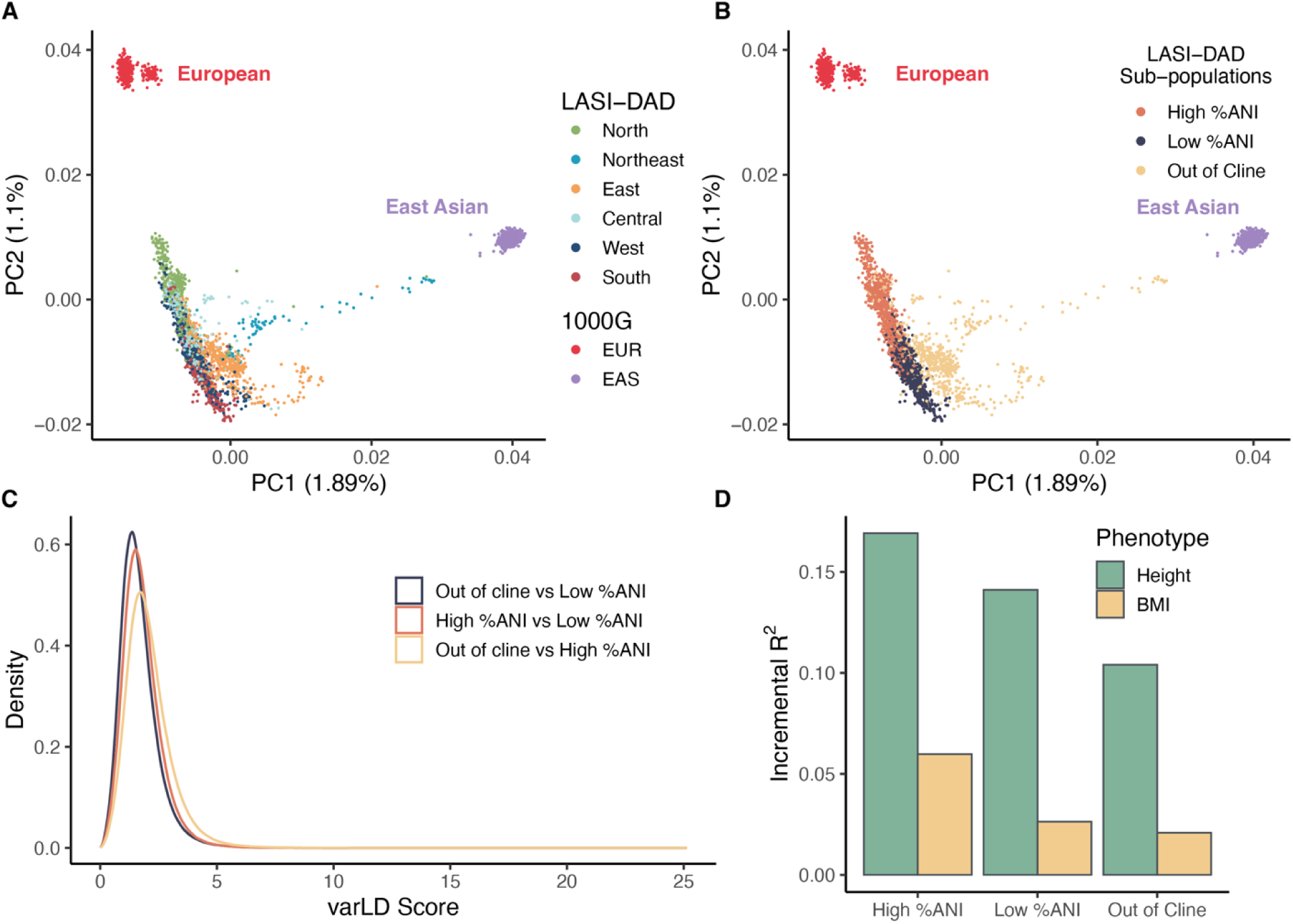
Linkage disequilibrium patterns in Indian sub-populations from LASI-DAD. Individuals from LASI-DAD were grouped into three sub-populations based on their proportion of Ancestral North Indian ancestry (%ANI) and presence on the North/South cline of India to facilitate Indian sub-population analysis. The three sub-populations included the high %ANI group (0.540 ≤ %ANI < 0.824; n = 935), low %ANI group (%ANI ≤ 0.540; n = 941), and the out of the North/South cline group (n = 731). (**A**) Principal component analysis (PCA) plot illustrating the genetic relationship among Indian, European, and East Asian populations. PCA was conducted using Indian samples from LASI-DAD along with European, South Asian, and East Asian samples from 1000G. The proportion of variance explained by each PC is indicated in the parentheses. LASI-DAD individuals are colored by their geographic region of birth in India to highlight the relationship between population structure and geography in India. (**B**) The same PCA plot with LASI-DAD individuals colored by their sub-population groups. (**C**) Distribution of variation in LD (varLD) scores evaluated between LASI-DAD sub-populations. (**D**) Predictive performance of polygenic risk scores (PRS) derived from GWAS of primarily European ancestry to the Indian sub-populations in LASI-DAD. Evaluated traits include height and body mass index (BMI), with GWAS summary statistics obtained from the GIANT consortium.

Despite overall modest differences in LD across LASI-DAD sub-populations, such differences can still impact the performance of various statistical analyses, such as the PRS. We evaluated the transferability of a height and body mass index (BMI) PRS derived from GWASs of primarily European ancestry to LASI-DAD individuals. We found that the predictive performance of PRS varied across LASI-DAD sub-populations (Figure 2D), with relative performance consistent with their genetic relatedness to the European population (Figures 2A-B). As expected, PRS performed the best in the high %ANI group due to its closer relatedness to the European population, achieving a prediction accuracy of 17% for height and 6% for BMI, as measured by incremental R^2^. In contrast, the prediction accuracy was 14% and 3% in the low %ANI group and 10% and 2% in the out-of-cline group for height and BMI, respectively. Despite the variation across sub-populations, the overall prediction accuracy in LASI-DAD remained substantially lower than that in the European population, where it reached as high as 44.7% for height^41^ and 10.2% for BMI.^42^ The limited transferability of PRS in LASI-DAD hinders its clinical utility, especially for South and East Indian populations who are more distantly related to the European population and benefit less from its large sample size. These findings suggest that more efforts need to be made toward genetic research in this unique population to reduce potential health disparities.^40^

### Providing LASI-DAD WGS data as an LD reference panel

We evaluated the utility of LASI-DAD WGS data as an LD reference panel to facilitate both LD lookup and various statistical analyses that rely on precise LD estimates. In the LD lookup panel, LASI-DAD included a total of 69.5 million variants, representing a 170% (43.8 million) and 213% (47.3 million) increase in variant coverage relative to the 1000G and TOP-LD panels, respectively (Figure 3A). To be included in the panels, these variants have at least one tagged variant with r^2^ ≥ 0.2. As expected, most variants in LASI-DAD that are not present in the other two panels have a relatively low MAF (e.g., < 1%) (Figure 3A). Interestingly, we found that the number of variants with a MAF ≥ 1% is smaller in the LASI-DAD panel (8.7 million) than in the 1000G (11.6 million) and TOP-LD (9.4 million) panels (Figure 3A). Presumably, this is because LASI-DAD represents a highly heterogenous yet nationally representative population that contains individuals with diverse histories of founder events.^38^ In contrast, the South Asian population in 1000G consists of individuals from only five sub-population groups, including 86 Bengali in Bangladesh; 103 Gujarati Indians in Houston, Texas; 102 Indian Telugu in the UK; 96 Punjabi in Lahore, Pakistan; and 102 Sri Lankan Tamil in the UK. Similarly, the only South Asian representation in TOPMed and its derived LD panel TOP-LD (n = 239) comes from the Pakistani Risk of Myocardial Infarction Study (PROMIS).^14, 52, 53^ The lack of genetic diversity in 1000G and TOP-LD for the South Asian population likely contributed to the larger number of variants with a relatively high MAF. The coverage of genetic variants remained the highest for the LASI-DAD panel regardless of the cutoff in r^2^ (Figure 3B). In addition, most variants in the LD lookup panel have at least one tagged variant with r^2^ ≥ 0.8, with proportions of variants being 82%, 76%, and 89% in the LASI-DAD, 1000G, and TOP-LD panels, respectively (Figure 3B).

**Figure 3.**
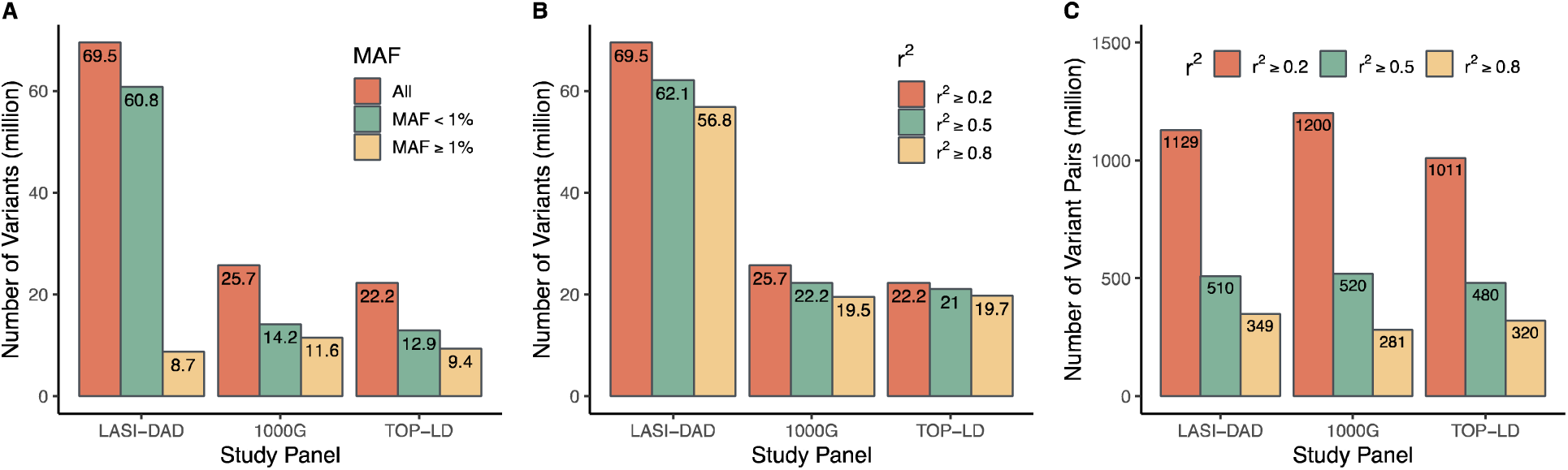
Number of autosomal variants included in different linkage disequilibrium (LD) reference panels. Three LD reference panels were compared, including Indian samples from LASI-DAD (N=2,607), South Asian samples from 1000G (N=489), and South Asian samples from TOPMed (N=239). (**A**) Number of variants stratified by the range of minor allele frequency (MAF). (**B**) Number of variants with at least one other variant in LD under different r^2^ cutoffs, where r^2^ represents the squared Pearson correlation coefficient between the phased haplotypes of a variant pair. (**C**) Number of variant pairs stratified by different r^2^ cutoffs.

Notably, the number of variant pairs remained largely similar across different LD lookup panels regardless of the cutoff in r^2^ (Figure 3C). Again, this is likely due to the lack of genetic diversity in 1000G and TOP-LD, resulting in each variant having more tagged variants in these two panels. Finally, we evaluated the consistency of LD estimates across the three lookup panels by examining the common set of variant pairs on chromosome 1. The Pearson correlation of r^2^ statistics was 0.94 between LASI-DAD and 1000G, 0.92 between LASI-DAD and TOP-LD, and 0.91 between 1000G and TOP-LD, indicating an overall high consistency across the three panels.

To demonstrate the advantage of LASI-DAD as an LD reference in statistical analyses, we conducted a PRS analysis where we compared the predictive performance of PRS constructed with either the LASI-DAD or 1000G SAS LD reference panel. To obtain PRS weights, we applied PRS-CS^26^ to four studies of South Asian ancestries, including a GWAS of height^41^ and an ExWAS of BMI^47^ from the GIANT consortium, and GWASs of both height and BMI from Genes & Health.^48^ We evaluated the prediction accuracy of PRS in LASI-DAD individuals. We found that the predictive performance of PRS was higher with the LASI-DAD panel than with the 1000G SAS panel for all four studies (Figure 4). Specifically, the LASI-DAD panel improved the prediction accuracy by 19.5% for height and 35.1% for BMI from the GIANT consortium, and by 2.2% for height and 2.1% for BMI from the Genes & Health. Notably, the improvement in PRS performance when using LASI-DAD as the reference panel was smaller for the Genes & Health data than for the GIANT data. This is likely because the LD patterns in LASI-DAD match more closely to those in GIANT than Genes & Health, as Genes & Health consists of British Pakistani and British Bangladeshi individuals, whereas a large proportion of individuals in GIANT are of Indian origin. These results highlight the benefits of LASI-DAD in serving as an LD reference panel.

**Figure 4.**
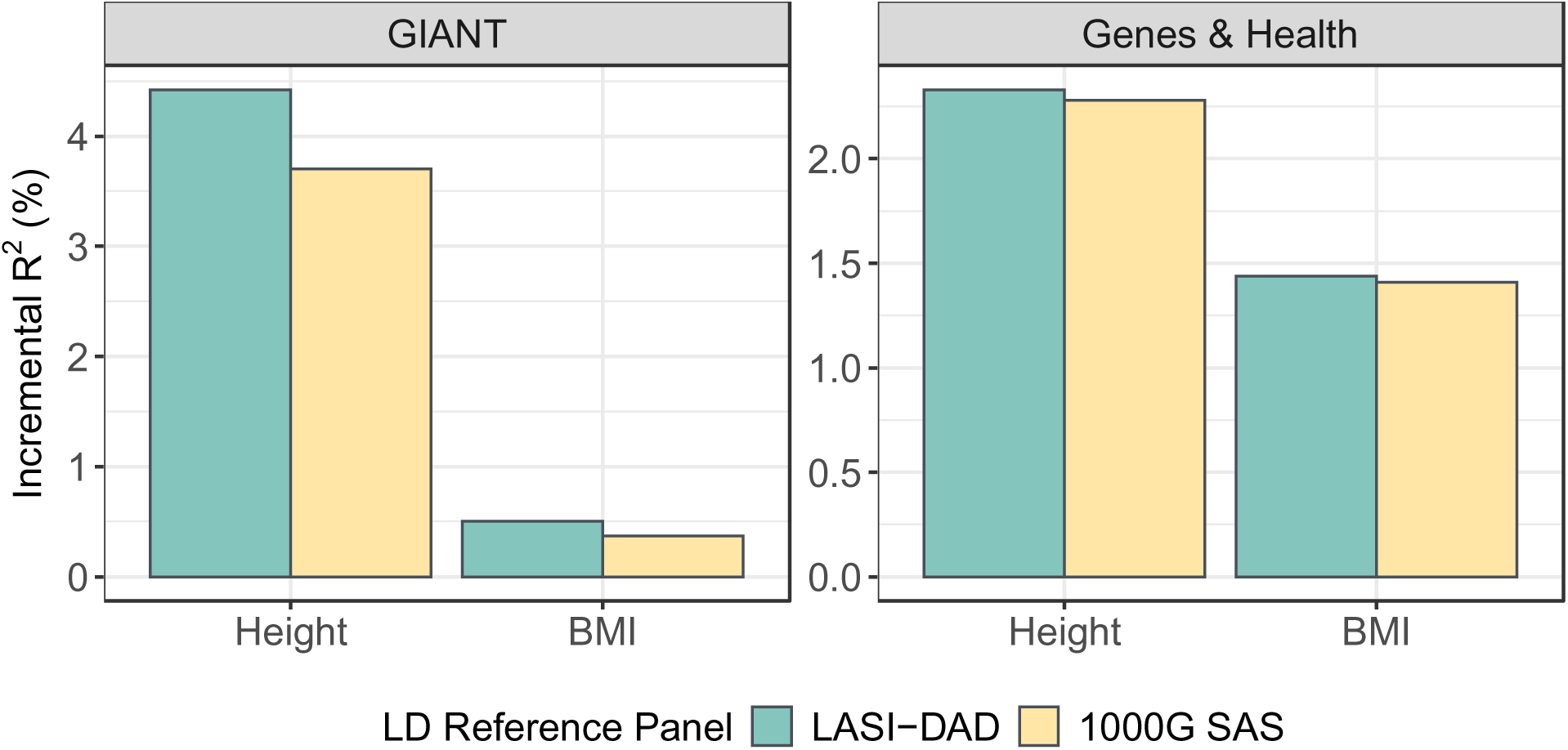
PRS analyses demonstrating the utility of LASI-DAD as an LD reference panel. The predictive performance of PRS constructed with the LASI-DAD and 1000G SAS LD reference panels were compared. Summary statistics for height and body mass index (BMI) were obtained from four studies of South Asian ancestry, including a GWAS of height and an ExWAS of BMI from the GIANT consortium, and GWASs of both height and BMI from Genes & Health. PRS weights were estimated by PRS-CS.

### Providing LASI-DAD WGS data as an imputation reference panel

We evaluated the utility of LASI-DAD WGS data as an imputation reference panel and compared it with the GAsP and TOPMed imputation panels. Specifically, we found that the LASI-DAD panel enabled the imputation of many Indian-specific variants that are not present in the other panels, especially those with a low MAF (Figure 5A). For rare variants with a MAF ≤ 1%, the LASI-DAD panel was able to impute a total of 61,843,011 variants, representing 182% (21,948,408) and 940% (5,943,775) increases relative to the TOPMed and GAsP panels, respectively. Similarly, for common variants with a MAF > 1%, the LASI-DAD panel was able to impute 8,783,797 variants, representing 4% (363,996) and 20% (1,449,768) increases relative to the TOPMed and GAsP panels. To ensure a fair comparison, we evaluated the imputation accuracy using the common set of variants that could be imputed by all three panels (Figure 5A).

**Figure 5.**
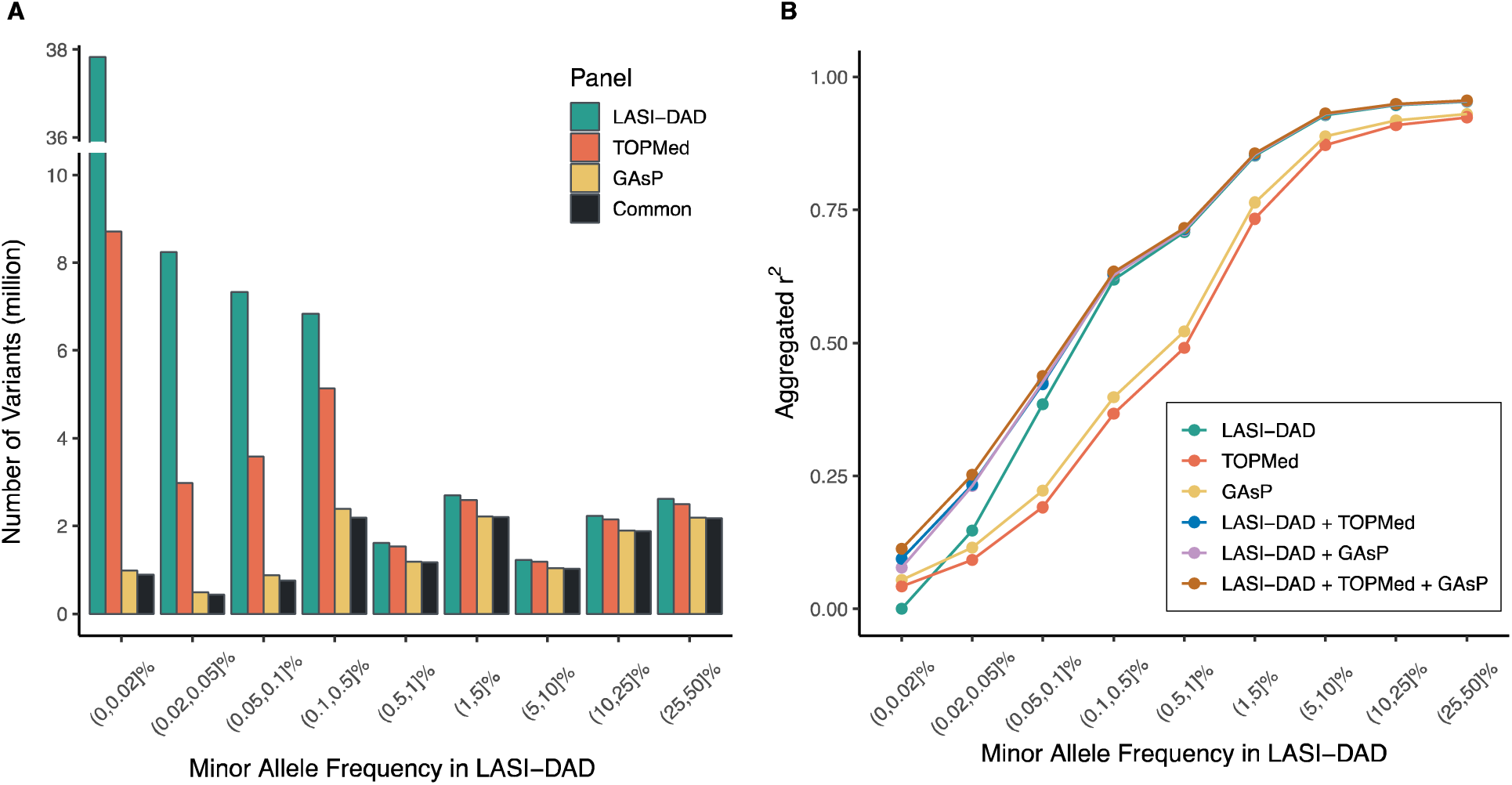
Evaluating LASI-DAD as a reference panel for genotype imputation. Compared reference panels include LASI-DAD, TOPMed, and GAsP. (**A**) Bar plots showing the number of genetic variants in LASI-DAD that can be imputed by each reference panel and by all three panels. (**B**) Imputation accuracy was evaluated for each reference panel, including meta-imputation to integrate the imputation results from either LASI-DAD and TOPMed, LASI-DAD and GAsP, or all three panels, across different minor allele frequency (MAF) ranges. The MAF was calculated within the full 2,680 LASI-DAD samples. To ensure a fair comparison, the imputation accuracy was evaluated using the common set of variants that can be imputed by all three panels. Aggregated r^2^ was used as the evaluation metric, calculated by grouping variants within a specified MAF range, stacking their imputed dosages and genotype calls from the sequencing data into two separate vectors, and then computing the squared correlation coefficient between the two vectors.

We found that the imputation with the LASI-DAD panel achieved substantially higher accuracy than with the other two panels for variants across most MAF ranges, except for singletons in the range of (0, 0.02]% (Figure 5B). For non-singletons, the LASI-DAD panel improved the imputation accuracy by 3% to 101% (mean = 38%) compared to the TOPMed panel and by 3% to 73% (mean = 27%) compared to the GAsP panel across different MAF ranges. The imputation performance is consistent with the sample size of South Asian individuals in each panel, with 1,654 samples in the GAsP panel and 644 in the TOPMed panel, both considerably smaller than that of the LASI-DAD panel. As expected, none of the panels can accurately impute singletons.

By combining imputation results from different reference panels, meta-imputation further improved the imputation accuracy, especially for variants with a low MAF (Figure 5B). Notably, meta-imputation using all three panels achieved the highest imputation accuracy across all MAF ranges. For variants with a relatively small MAF (≤ 0.1%), meta-imputation increased the accuracy by 14% to 107% (mean = 64%) compared to the best-performing reference panel. On the other hand, for variants with a relatively large MAF (> 0.1%), incorporating additional reference panels beyond LASI-DAD offered only limited benefits, with imputation accuracy improving by 0.2% to 2% (mean = 0.8%) compared to the LASI-DAD panel. These results highlight the advantages of meta-imputation, in complementing the LASI-DAD panel, for imputing relatively rare variants in the Indian population.

Finally, we found that the imputation accuracy varied for samples from different states or territories of India, with the LASI-DAD panel exhibiting the smallest variation (Figure S3). This variation in imputation accuracy can be explained to some extent by the sample composition of the reference panels. For example, the TOPMed panel, which consists nearly 50% European samples, exhibits a strong correlation between the imputation accuracy and the average %ANI for each state (R^2^ = 0.77; Figure S4). This is consistent with the fact that samples with higher %ANI have closer relatedness to West Eurasians. In contrast, the correlation is much weaker with the LASI-DAD (R^2^ = 0.42) or GAsP (R^2^ = 0.15) panel. These results highlight the importance of having a large and nationally representative dataset for the Indian population to achieve the best imputation performance due to its complex population structure.

## Discussion

As the largest and most nationally representative WGS study of the Indian population, LASI-DAD provides a unique opportunity to serve as a valuable resource for both LD and genotype imputation in this population. In the present study, we characterized the LD patterns in LASI-DAD by identifying LD blocks and evaluating regional differences in LD between the Indian population from LASI-DAD and four super-populations from the 1000G. Besides super-populations, we explored the population structure of India by evaluating LD patterns across Indian sub-populations. We evaluated the utility of LASI-DAD WGS data as an LD reference panel to facilitate both LD lookup and various statistical analyses that rely on precise LD estimates. Finally, we evaluated the utility of LASI-DAD as an imputation reference panel. As an LD reference panel, LASI-DAD provides a more comprehensive representation of genetic variation/structure in India and facilitates downstream LD-dependent analyses. As an imputation panel, it improves imputation accuracy compared to existing panels and demonstrates more robust performance across different Indian states or territories.

The comparison of LD patterns between LASI-DAD and four super-populations from 1000G confirms that the African population has the largest number of LD blocks and smallest overall block sizes. In comparison, the European, East Asian and South Asian populations, which all migrated from Africa, have a comparable number of blocks with similar block sizes. As expected, LASI-DAD has a similar LD structure as the 1000G South Asian population, in terms of both the number and size of LD blocks, according to either the broad-scale LD blocks identified by LDetect or the fine-scale LD blocks identified by BigLD. However, our analysis still reveals some regions with considerable LD differences between LASI-DAD and 1000G SAS. This heterogeneity is further highlighted by global genetic principal components, showing that LASI-DAD significantly broadens the genetic boundaries typically seen in the 1000G South Asian panel (Figure S5). Moreover, due to a significant increase in sample size, LASI-DAD interrogates significantly more (170%) variants, likely specific to the Indian/South Asian population, that are not observed in the 1000G SAS panel. Given the improved representation and coverage, it is not surprising that the LASI-DAD panel, in comparison to 1000G SAS panel, improves polygenic score construction and prediction in South Asians. We expect similar improvement in other LD-based analyses that target Indian and South Asian populations.

Unfortunately, the sample sizes of South Asian GWASs are limited; thus, the overall prediction accuracy of the corresponding PRS is much lower than that based on the European GWASs (Figure 2D versus Figure 4). The next important step is to expand GWAS efforts in South Asian populations, for which a good imputation reference panel is needed to generate high quality genotype data. LASI-DAD can well serve that purpose as it improves imputation accuracy over existing popular reference panels for South Asians, including TOPMed and GAsP. Importantly, it has more robust performance across all Indian states or territories, likely due to its unbiased representation of the Indian population. Our analysis further highlighted the potential advantage of leveraging multiple reference panels via meta-imputation to further improve imputation accuracy. All of these demonstrate that LASI-DAD is a valuable genetic reference panel for Indian and South Asian populations.

The Indian population is diverse and shaped by multiple ancestral influences. Specifically, the majority of Indians are part of a North/South cline and are admixed with varying degrees of ANI (related to European, Central Asia and Middle Eastern populations) and ASI (related to endogenous ancestry). Meanwhile, a small proportion of Indians who are in the East have an additional ancestral influence from East Asians. Our analysis by LASI-DAD sub-population confirms that this population structure leads to LD differences among sub-populations, with the largest difference observed between North Indians (high %ANI group) and East Indians (out-of-cline group), while South Indians (low %ANI group) are positioned in between. Since North Indians are most closely related to Europeans, the predictive performance of European-based PRSs shows a similar gradient across these sub-populations, with highest performance in North Indians, followed by South Indians, and then East Indians. This is consistent with many studies, showing that the transferability of PRS decreases as the genetic distance between target population and discovery population increases.^54^ The heterogeneity could also affect imputation accuracy across sub-populations when the reference panel is biased. This heterogeneity and potential consequence on health inequity underscore the importance of including diverse groups and having population-representative Indian samples, as exemplified by LASI-DAD.

While LASI-DAD has proven to be a valuable resource for serving as a reference panel, several opportunities are possible to further improve its utility. First, while LASI-DAD represents the largest and most nationally representative WGS study, it remains insufficient to fully capture the extensive genetic diversity of the Indian population, which includes over 4,500 anthropologically defined groups. Substantial efforts to collect more samples are still needed to achieve a more comprehensive representation. Second, most existing genetic studies analyze South Asian populations collectively as a single group. However, South Asia is a region that spans eight countries, making up about 25% of the world’s population and having a rich genetic diversity.^52^ As a result, LD patterns observed in one South Asian population may not accurately reflect those in another. This partly explains why the improvement in PRS performance when using LASI-DAD as the reference panel was smaller for the Genes & Health data than for the GIANT data, as the Genes & Health data consists of British Pakistani and British Bangladeshi, whereas a large proportion of individuals in the GIANT data are of Indian origin. To achieve the best results in statistical analyses with GWAS summary statistics, identifying the most appropriate reference panel with LD patterns that best match the GWAS population is critical. In line with this, an adaptive approach that combines multiple reference panels, akin to the meta-imputation approach, may provide an alternative way for improving the LD reference panel. However, this would require the development of new statistical frameworks or analytical procedures.

In summary, we have presented LASI-DAD, the largest and most nationally representative WGS data from India, as a reference panel for LD and genotype imputation. This valuable resource fills a critical gap in the field of genetics and could greatly facilitate various genetic analyses in Indians and South Asians. Meanwhile, the genetic diversity demonstrated in this study also underscores a critical need to expand data collection and enhance representativeness in this population.

## Acknowledgements

We thank the LASI-DAD participants for being part of the research project. We thank Dr. Priya Moorjani at the University of California, Berkeley for her guidance on defining the LASI-DAD sub-populations. This study was supported by the National Institute on Aging (R01 AG051125, U01 AG064948, and U54-AG-052427). The funders had no role in study design, data collection and analysis, decision to publish or preparation of the manuscript.

## Author contributions

W.Z., J.S., and S.K. conceived the idea. J.L. and S.K. provided funding support. S.D., A.B.D., and J.L. led data collection in LASI-DAD. Y.Y.L, G.D.S and L.-S.W. processed and cleaned the whole genome sequencing data. Z.L. and W.Z. designed the experiments. Z.L. conducted the data analyses. Z.L., W.Z., and J.S. wrote the manuscript with input from X.Z., Y.Y.L, and G.D.S. All authors read and approved the final manuscript.

## Institutional review board statement

Investigations were conducted in accordance with the principles outlined in the Declaration of Helsinki (1975, revised in 2013). The study was approved by Institutional Review Boards at the University of Southern California (UP-15-00684) and the University of Michigan (HUM00166956).

## Informed consent statement

Written informed consent was obtained from all participants.

## Data availability statement

Whole genome sequencing data for the Diagnostic Assessment of Dementia for the Longitudinal Aging Study of India (LASI-DAD) is available from the National Institute on Aging Genetics of Alzheimer’s Disease Data Storage Site (NIAGADS), accession number: NG00067 – ADSP Umbrella. Phenotype data is available at the Gateway to Global Aging website, https://g2aging.org/. The LASI-DAD LD reference panel is available at https://github.com/zhengli09/LASI_DAD-LD-Reference-Panel. We are currently working with the Michigan Imputation Server (https://imputationserver.sph.umich.edu) to host the LASI-DAD imputation panel; however, it is not yet available. Researchers wishing to access the imputation panel should contact the corresponding authors.

