## Supplemental Tables and Figures for "A reference panel for linkage disequilibrium and genotype imputation using whole-genome sequencing data from 2,680 participants across India" (from another version of this preprint: version 1, 2025-07-04)

**Table S1. Sample sizes of Indian states and union territories by geographic region in LASI-DAD.**

| Geographic region of India | State/Territory | Sample size |
|---|---|---|
| North | Jammu & Kashmir | 90 |
| | Punjab | 112 |
| | Haryana | 171 |
| | Delhi | 120 |
| | Rajasthan | 151 |
| Northeast | Assam | 72 |
| East | Bihar | 137 |
| | West Bengal | 187 |
| | Odisha | 216 |
| Central | Uttarakhand | 68 |
| | Uttar Pradesh | 154 |
| | Madhya Pradesh | 77 |
| West | Gujarat | 228 |
| | Maharashtra | 185 |
| South | Telangana | 171 |
| | Karnataka | 122 |
| | Tamil Nadu | 159 |
| | Kerala | 260 |

**Table S2. Summary of linkage disequilibrium blocks as identified by BigLD.**

| Data Source | Population | Number of Blocks | Block Length (kb)* | | Number of SNPs^ | |
|---|---|---|---|---|---|---|
| | | | Median | Mean | Median | Mean |
| LASI-DAD | Indian | 106,383 | 5.9 | 23.5 | 20 | 74 |
| 1000G | African | 143,383 | 3.1 | 17.9 | 20 | 103 |
| | European | 98,465 | 6.1 | 25.9 | 22 | 87 |
| | South Asian | 108,751 | 5.4 | 23.4 | 21 | 83 |
| | East Asian | 96,415 | 6.8 | 26.2 | 22 | 79 |

*SNPs with a MAF ≥ 5% were used for block detection.
^The number of SNPs was counted across all SNPs with MAF ≥ 1%.

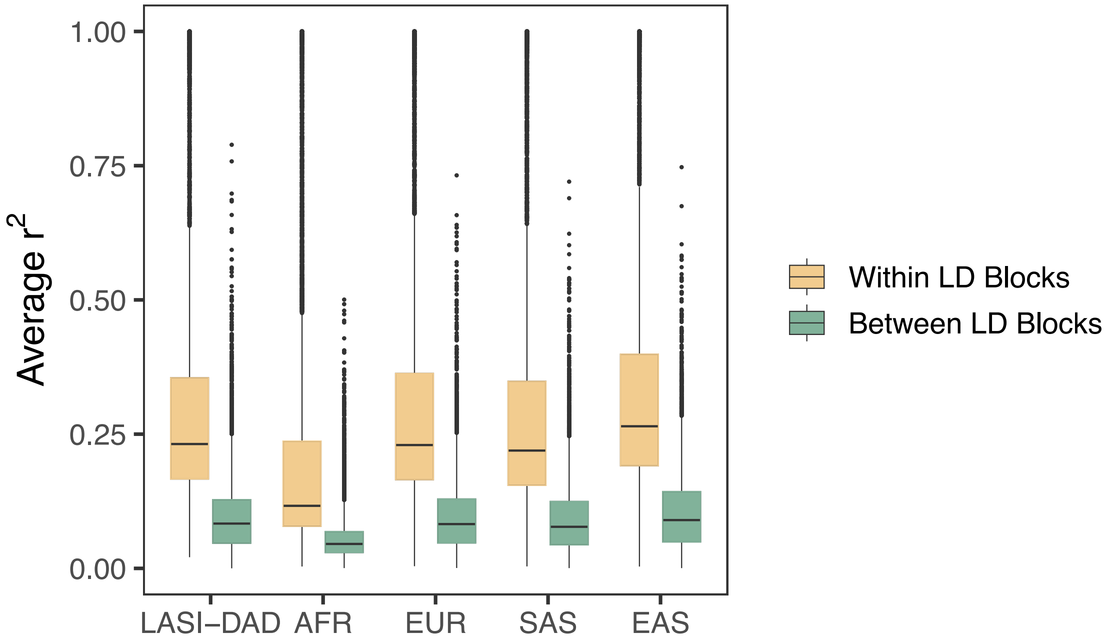


**Figure S1. Distribution of average linkage disequilibrium (LD) on chromosome 1.** The average LD was evaluated either between all pairs of SNPs within each LD block or between SNPs in each LD block and its adjacent two blocks. LD blocks were identified by BigLD at fine scales. LD was evaluated as the squared correlation coefficient ($r^2$) of genotypes between a SNP pair.

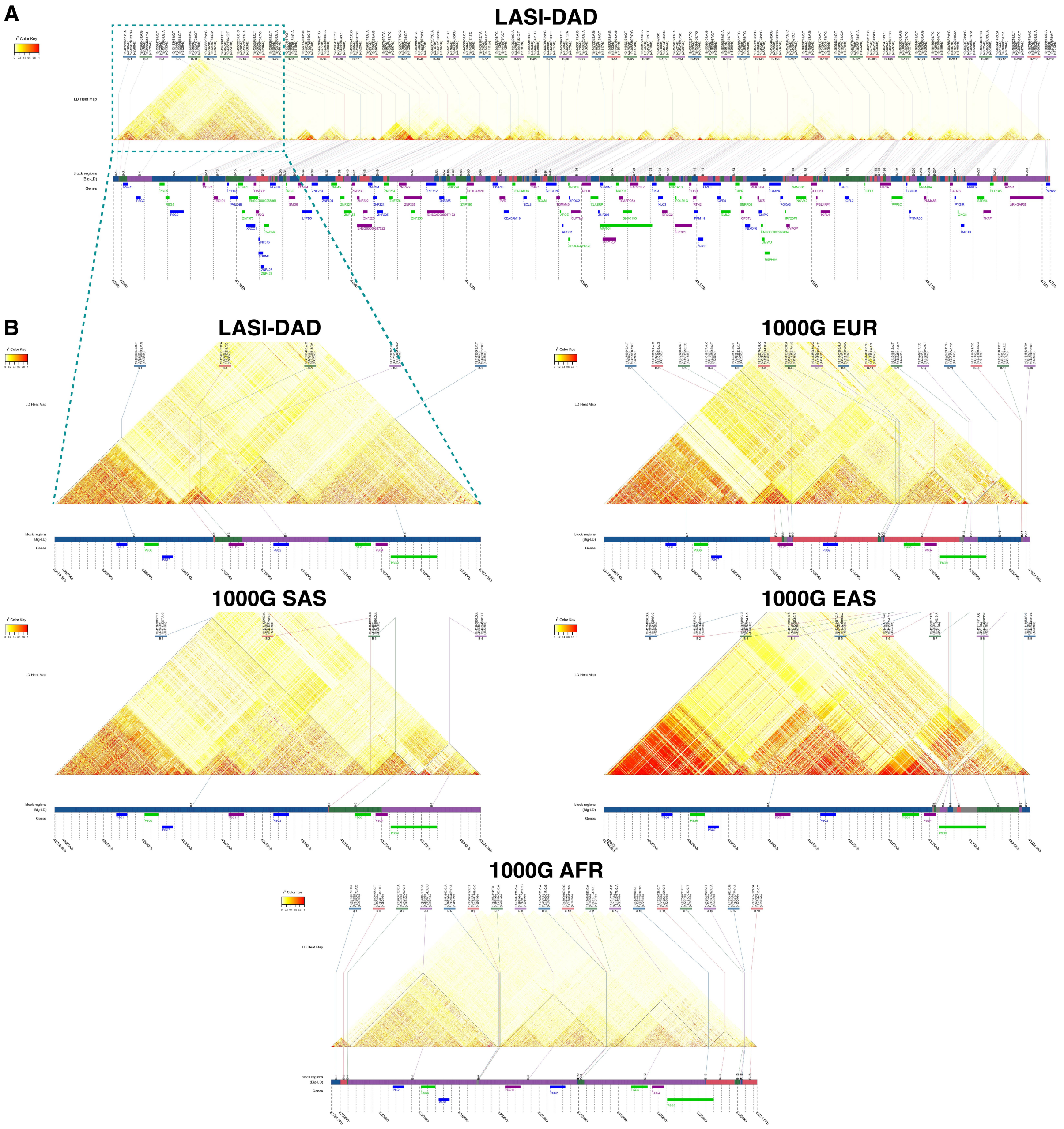


**Figure S2. Linkage disequilibrium (LD) patterns of an example genomic region.** (**A**) Visualization of an example LD block identified by LDetect from LASI-DAD. The LD block spans the region from 42,965,392 to 47,034,923 on chromosome 19. The fine-scale LD blocks identified by BigLD and their boundary SNPs are indicated at the top, while genes are indicated at the bottom. (**B**) Zoom-in of a particular region with extensive variation in local LD structures across populations, including Indian from LASI-DAD, and African (AFR), European (EUR), South Asian (SAS), and East Asian (EAS) from the 1000 Genomes Project (1000G).

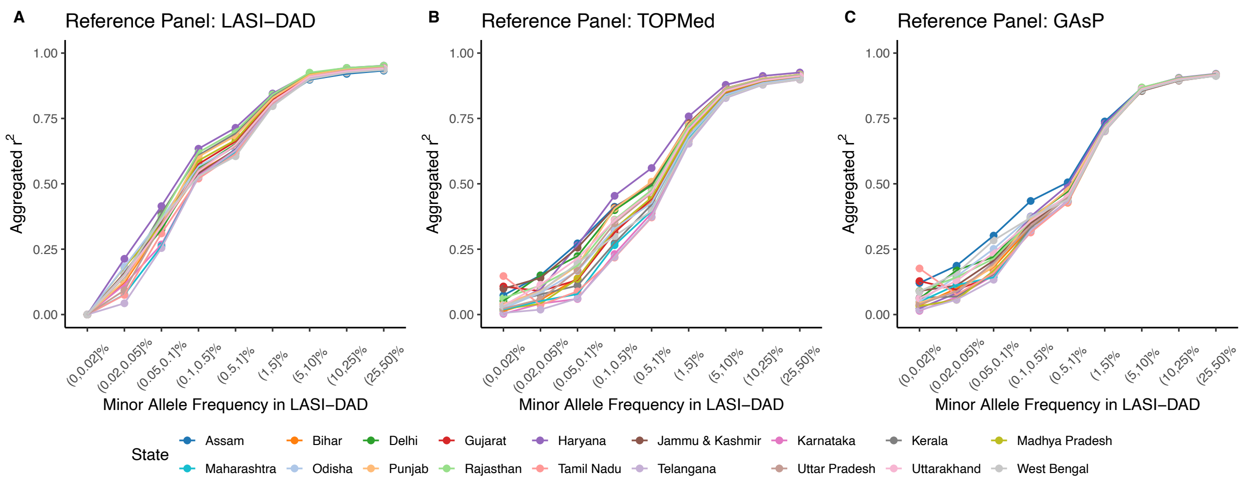


**Figure S3. Imputation accuracy by Indian state/territory**. Imputation accuracy was evaluated for samples from each Indian state or union territory across different minor allele frequency (MAF) ranges. Compared imputation reference panels include LASI-DAD, TOPMed, and GAsP. The MAF was calculated within the full 2,680 LASI-DAD samples. To ensure a fair comparison, the imputation accuracy was evaluated using the common set of variants that can be imputed by all three panels. Aggregated $r^2$ was used as the evaluation metric, calculated by grouping variants within a specified MAF range, stacking their imputed dosages and genotype calls from the sequencing data into two separate vectors, and then computing the squared correlation coefficient between the two vectors.

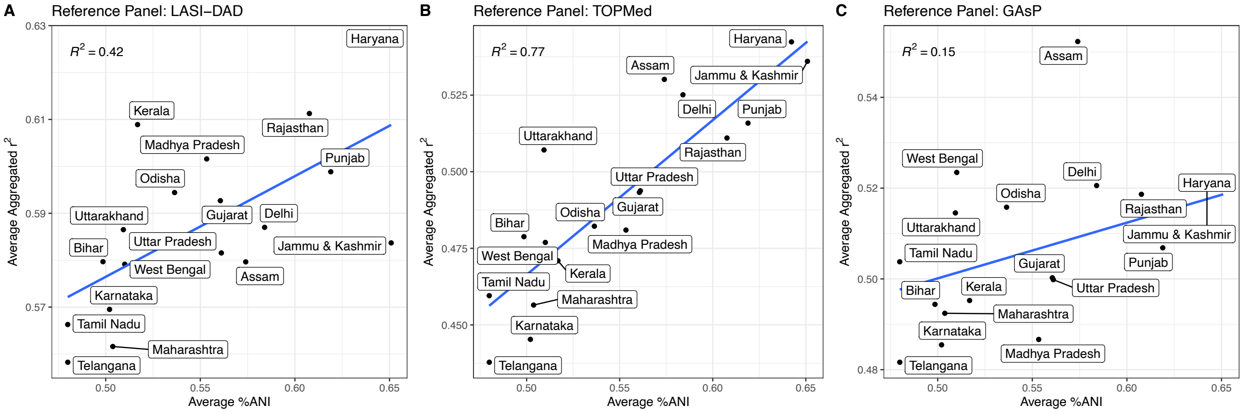


**Figure S4. Average proportion of Ancestral North Indian (%ANI) in each state/territory of India versus average aggregated imputation $r^2$ with different panels.** The aggregated $r^2$ was computed with imputed dosages from panel (**A**) LASI-DAD, (**B**) TOPMed, or (**C**) GAsP.

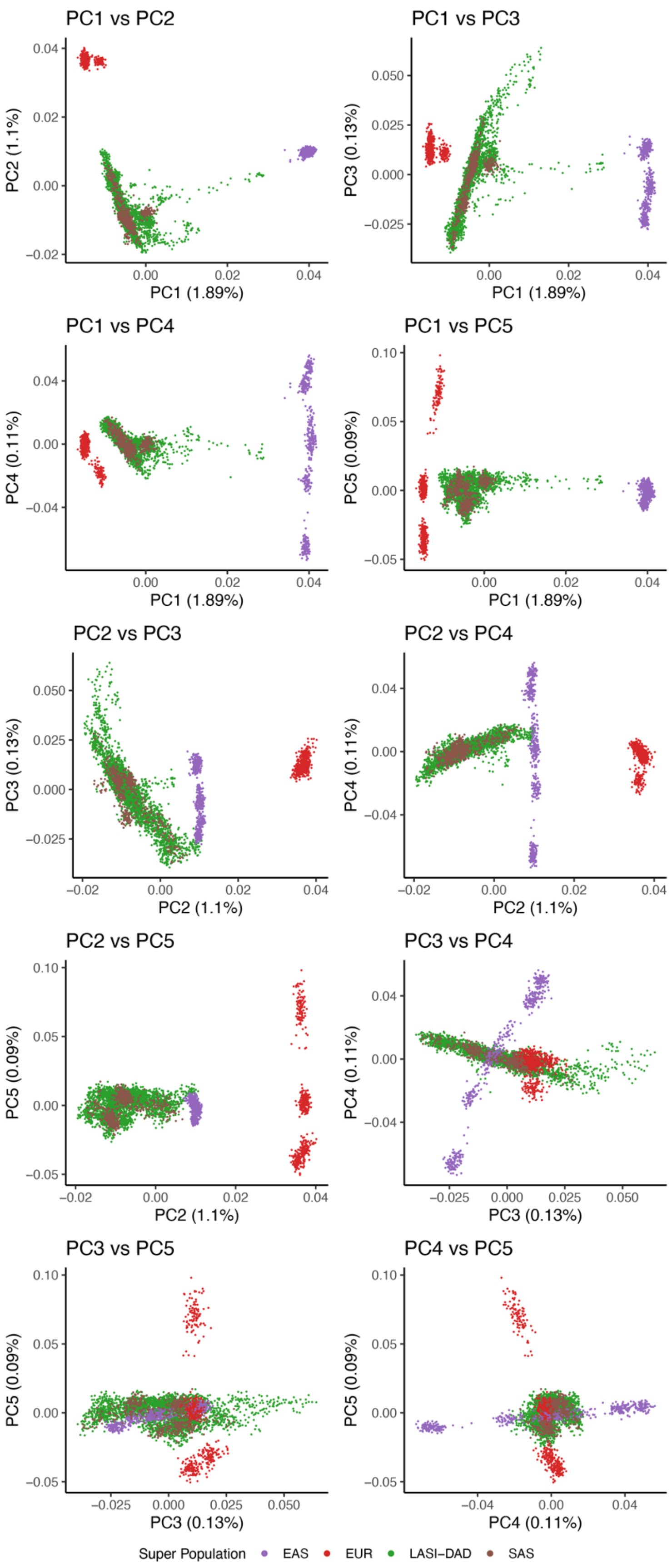
PC1 vs PC2
PC1 vs PC3
PC1 vs PC4
PC1 vs PC5
PC2 vs PC3
PC2 vs PC4
PC2 vs PC5
PC3 vs PC4
PC3 vs PC5
PC4 vs PC5
PC1 (1.89%)
PC2 (1.1%)
PC3 (0.13%)
PC4 (0.11%)
PC5 (0.09%)
Super Population
EAS
EUR
LASI−DAD
SAS

**Figure S5. Principal component analysis (PCA) plots.** PCA was conducted using Indian samples from LASI-DAD along with European, South Asian, and East Asian samples from 1000G. The proportion of variance explained by each PC is indicated in the parentheses.
